# The MRC IEU OpenGWAS data infrastructure

**DOI:** 10.1101/2020.08.10.244293

**Authors:** Ben Elsworth, Matthew Lyon, Tessa Alexander, Yi Liu, Peter Matthews, Jon Hallett, Phil Bates, Tom Palmer, Valeriia Haberland, George Davey Smith, Jie Zheng, Philip Haycock, Tom R Gaunt, Gibran Hemani

**Author notes:** Equal contributorship (Ben Elsworth, Matthew Lyon, Tom R Gaunt, Gibran Hemani).

## Abstract

Data generated by genome-wide association studies (GWAS) are growing fast with the linkage of biobank samples to health records, and expanding capture of high-dimensional molecular phenotypes. However the utility of these efforts can only be fully realised if their complete results are collected from their heterogeneous sources and formats, harmonised and made programmatically accessible.

Here we present the OpenGWAS database, an open source, open access, scalable and high-performance cloud-based data infrastructure that imports and publishes complete GWAS summary datasets and metadata for the scientific community. Our import pipeline harmonises these datasets against dbSNP and the human genome reference sequence, generates summary reports and standardises the format of results and metadata. Users can access the data via a website, an application programming interface, R and Python packages, and also as downloadable files that can be rapidly queried in high performance computing environments.

OpenGWAS currently contains 126 billion genetic associations from 14,582 complete GWAS datasets representing a range of different human phenotypes and disease outcomes across different populations. We developed R and Python packages to serve as conduits between these GWAS data sources and a range of available analytical tools, enabling Mendelian randomization, genetic colocalisation analysis, fine mapping, genetic correlation and locus visualisation.

OpenGWAS is freely accessible at https://gwas.mrcieu.ac.uk, and has been designed to facilitate integration with third party analytical tools.

## Introduction

Genome-wide association studies (GWAS) have contributed to our understanding of the biology of many complex traits and diseases ^1^. Since their emergence nearly 15 years ago many thousands of GWAS studies have been published, and the results from these studies offer a portal into understanding disease biology, complex trait architecture and evolutionary history. This paper describes our efforts to harmonise results of these studies into a platform that is freely programmatically accessible for a range of analytical tools.

Genetic variation has two important properties. First, it represents quasi-random perturbations of genomic regions^2^. When tested for association with a trait, genetic variants can be used to approximate the counterfactual effect of a variant substitution on a trait^3^. Second, due to imputation and a focus on common variants, the genetic variants analysed in GWAS are finite in number, with a large proportion of variants consistently included across all GWAS^4^. These two properties make GWAS results a powerful tool for understanding the shared genetic properties of different traits, without the need for individual-level genetic data^5^. Methodological advancements have been made in particular in Mendelian randomization (MR) ^6^, genetic colocalisation ^7–9^ and genetic correlation ^10,11^ that require only GWAS summary data to explore the shared aetiology of two or more traits.

While the rate of GWAS summary data generation is accelerating, the collective value of these resources do not meet their full potential if they are not made publicly available, or are only made available in non-standard formats and dispersed across the internet. In 2015 we launched the MR-Base platform which systematically collected complete GWAS summary data into a single database ^12^. The database was accessible through the TwoSampleMR R package ^13^, which focused on querying the data to automate MR analyses ^6^. Since then several other similar resources have appeared ^14,15^, and development of the MR-Base platform has also continued.

Now superseding the original MR-Base database, the MRC IEU OpenGWAS database aims to be more open, scalable and accessible for other analytical approaches and third-party software. This paper describes the resource, and explains how and why various design choices were made. In the Discussion we detail the practical and technical barriers that remain in building a homogeneous resource of GWAS summary data.

## Results

The OpenGWAS infrastructure incorporates four domains - (1) *data are sourced and imported* before being (2) *processed for format and quality control* in order to be (3) *served publicly* and (4) *connected to a range of analytical tools* (Figure 1).

**Figure 1:**
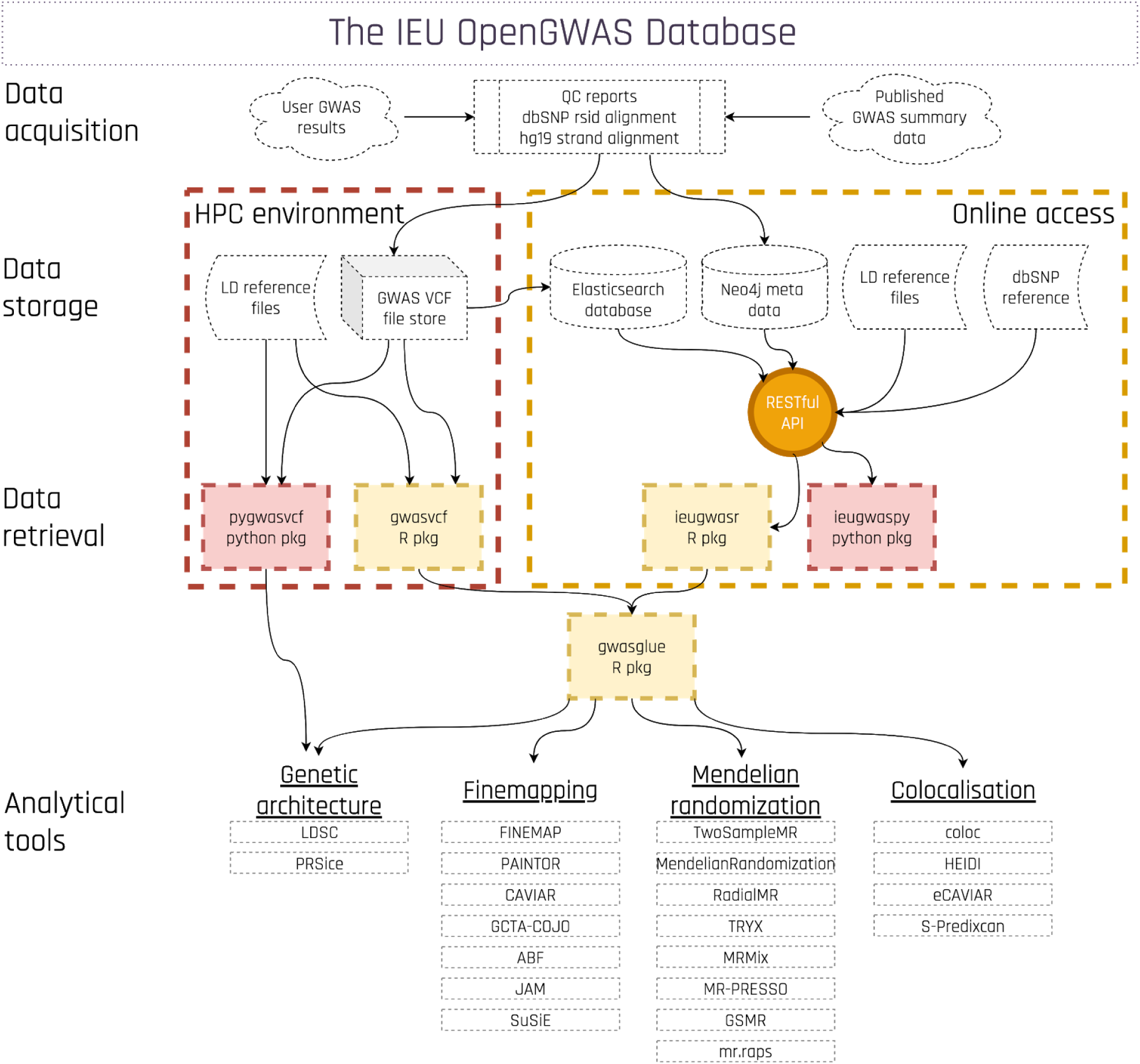
A schematic of the GWAS summary data infrastructure that constitutes the IEU OpenGWAS database. Each of the data retrieval methods are open to be linked to other third party tools.

### Summary of the database contents

At the time of writing there are 14582 complete and publicly available GWAS summary datasets within OpenGWAS, where ‘complete’ refers to all analysed variants being retained regardless of p-value. This corresponds to a total of 125.8 billion genetic associations. The data are typically sourced from results made public by specific GWAS consortia (representing 196 unique publications) or those that were analysed in bulk on biobanks (such as UK Biobank and Biobank Japan). At this point the data are predominantly from samples of European ancestry, but as new biobanks around the world begin publishing their GWAS results the populations represented in the database will increase in diversity.

Included amongst the datasets are a wide range of traits. At the time of writing there are 4126 binary traits, 725 metabolites, 3371 proteins, 3143 brain imaging phenotypes, and 3217 other continuous phenotypes. In addition to the complete GWAS summary data, we have also extracted the independent top hits for every dataset, totalling 116918 independent signals in which 7109 datasets have at least one hit. All publicly available datasets can be searched and browsed online or through various programming interfaces as we go on to describe^16^.

### Data harmonisation and quality control

Datasets obtained from different sources can be difficult to analyse jointly. Even after imputation, GWAS results can vary in terms of how variant identifiers are presented, which alleles are used to represent the effect allele, which version of dbSNP is used to name the variants, etc. We developed a harmonisation pipeline (https://github.com/MRCIEU/gwas2vcf) that aligns the non-effect allele for every variant to the human genome reference sequence, trims alternative alleles to their minimal representation ^17^ and matches the variant to a consistent build of dbSNP ^18^ (Figure 2).

**Figure 2.**
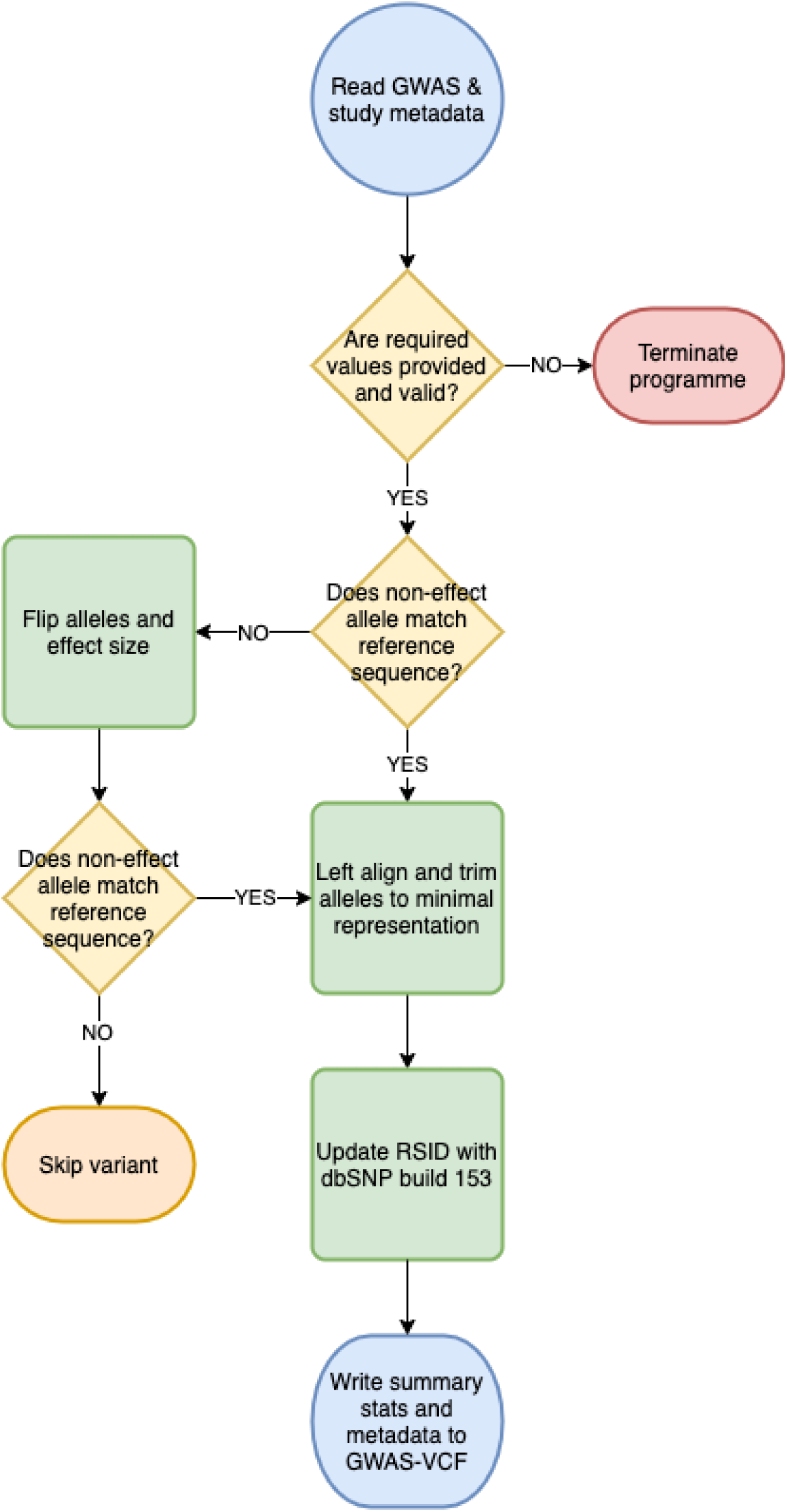
GWAS harmonisation flowchart.

In addition, we generate and publish a report for every dataset that evaluates the sanity of the reported statistics. The report includes a summary of the estimated heritability, genomic inflation and genetic confounding of the dataset; a list of independent sites after strict LD-based clumping; a comparison of allele frequencies against 1000 genomes reference data ^19^; and a comparison of Z-scores estimated from the reported effect sizes and standard errors against those calculated from the reported p-values.

### Online access to the database

The primary goal of creating a comprehensive database of harmonised GWAS summary statistics is to enable easy flow of that data through to analytical tools. There are two stages to this process: (1) fast and easy querying of the data with different programming languages; (2) converting the query results into formats ready to be analysed by different tools.

Access to the database is mediated by a representational state transfer (RESTful) application programming interface (API) ^20^, which allows positional queries based on dbSNP rsid, chromosome:position, or a genomic range. Where a requested rsID is not available for a particular dataset, it will return the result for a proxy variant (a variant in high linkage disequilibrium, LD, with the requested variant) (with aligned alleles) instead. Queries by p-value are also available, returning all variants with association p-values below that p-value, or (optionally) a subset of these based on automated, on-the-fly clumping. The API provides options to generate LD matrices for a given set of variants and perform LD clumping on a given set of variants. The API also makes it possible to retrieve the effect of a single variant or chromosome-position range on all traits in the database (i.e. a phenome-wide association study, PheWAS).

For convenience, all the API endpoints can be accessed directly (e.g. using curl) or through either an R package (ieugwasr ^21^) or a python package (ieugwaspy ^22^).

### Accessing the data for high performance computing

If a high volume of queries needs to be performed, or the results of a query will typically be very large, performing the queries through the API will be sub-optimal both for the user and for the OpenGWAS infrastructure, as intensive, automated queries run in parallel can influence the performance of the database. To this end we have made the complete data available already harmonised for download as flat files that can be utilised in any high performance computing (HPC) environment. Each GWAS summary dataset is available for download in GWAS-VCF format ^23^, designed to optimise data fidelity and query speed relative to standard tabular text files. We provide R (gwasvcf ^24^ [23]) and python (pygwasvcf ^25^) packages that can perform rapid queries on these file formats, therefore simplifying parallel processing of many large queries.

### Connecting the queried data to analytical tools

Both the gwasvcf ^24^ and the ieugwasr ^26^ R packages retrieve GWAS summary data into set data structures within R, while the ieugwaspy ^22^ package retrieves GWAS summary data into Python dictionaries (easily convertible to Pandas dataFrames). Across the discipline, many different software tools have been developed for MR, genetic colocalisation analysis and fine-mapping analysis. Each requires slightly different data fields and formats, may require LD matrices for the set of variants under analysis, and may require multiple datasets to be harmonised for joint analysis. To this end we developed the gwasglue R package ^27^. This is a repository of R methods that connect either the API data or GWAS-VCF data directly to a broad set of analytical tools (Figure 1). As new software applications become available that are relevant to GWAS summary data, the gwasglue package will be updated to accommodate the necessary data structures to enable connectivity between the data and the analytical methods.

### Future data harvesting

Future data harvesting will be from multiple sources. Some GWAS studies will perform detailed analysis of one or few traits, which can be uploaded through an API. Other GWAS studies will analyse hundreds or thousands of traits, which we process using a HPC pipeline. Any new complete GWAS summary datasets that have been harmonised by the EBI-NHGRI GWAS catalog ^15^ are automatically integrated into OpenGWAS. We strongly encourage users and researchers performing GWAS to upload their published summary data to the EBI-NHGRI GWAS catalog, which will then be automatically mirrored in OpenGWAS for more extensive analytical options. Unpublished GWAS summary datasets (that are not eligible for inclusion in the EBI-NHGRI GWAS catalog) can be submitted directly to the OpenGWAS database using the request system here: https://github.com/mrcieu/opengwas-requests.

We note also that in some instances users request that uploaded datasets are embargoed for a specified time period. OpenGWAS allows user-specific access to specified embargoed datasets with access being authenticated using the Google OAuth2.0 protocol.

## Discussion

We have created a resource that harmonises and shares the results of a diverse array of published and unpublished GWAS. This is the largest resource of its kind described to date, and has been designed to accelerate the joint analysis of multiple traits in a free and open manner. The data can be downloaded or queried directly through an API, and we have made available tools in R and Python to connect to the resource. Users have options to operate on the data through the cloud or through high performance computing infrastructures. We also developed an R package that automates the channelling of the data in OpenGWAS to a diverse set of analytical tools. The automated data import pipelines that we developed enables future growth of the database and provides transparency over the data processing steps through the publication of QC reports for each dataset. A comparison against other similar resources is shown in Table <u>4</u>.

We do emphasise, however that there are a number of complex features and practices of GWAS in this diverse and evolving field. Hence the analysis of summary statistics (where users have no control over sample selection, individual-level data QC or statistical models) raises some important considerations for users of OpenGWAS

### Population differences between studies

While most datasets in OpenGWAS derive from European ancestry samples, there are substantial population differences between and within country ^28,29^. This has implications for LD structure, assumptions of panmixia and population structure. Bias can arise in methods such as two sample MR and genetic colocalisation when two datasets are analysed jointly but with different underlying LD structures. If two datasets are analysed jointly it often assumes panmixia - i.e. that the underlying causal structure is identical across the two samples. This could further be violated if two studies include individuals from different environments or from different time points. Similarly, population structure could introduce apparent genetic differences between studies if they have varying ancestral backgrounds.

### GWAS methods vary between studies

Various statistical methods are in common use for performing GWAS. Linear models and linear mixed models can have different properties, especially with regards to how controls for population structure influence the estimated effect sizes ^30^. LD score regression may give substantially attenuated results for GWAS summary data generated using LMM ^11^. Studies obtained from family data (e.g. transmission disequilibrium test results from trios) could differ substantially from population based estimates as they may more adequately control for population structure as well as family structure, such as assortative mating or dynastic effects ^2,31,32^.

### Controlling for covariates differs between studies

On the assumption that covariates included in a model are a cause of the target trait, a number of differences between studies can arise if the set of covariates used differ. For example, a GWAS of type 2 diabetes that has controlled for adiposity will have a different genetic profile than one that does not, because adiposity is causal for type 2 diabetes, and the genetic effects that influence type 2 diabetes through adiposity will be attenuated ^33^. If more covariates are controlled for in one study compared to another, and the residuals have not been scaled, effect size estimates should not be altered drastically. However, the variance explained by the variants will appear inflated if there are more covariates controlled for in one study.

On the assumption that the covariate included in a model is itself caused by the target trait, complications could potentially arise due to collider bias. Here, genetic influence on the covariate could become associated with the trait, leading to spurious associations or biased genome-wide effect estimates ^34^. Such a situation is difficult to guard against, so where possible the trait metadata in OpenGWAS includes information about covariates adjusted for in the model.

### Quality control decisions differ between studies

Different GWAS consortia and biobank datasets have varying approaches to performing quality control. In particular, decisions about imputation accuracy thresholds can introduce heterogeneity between studies. If there are differences in imputation accuracy between two studies then the one with poorer imputation will have stronger attenuation of effect size estimates, which could introduce heterogeneity in MR studies, loss of power in colocalisation studies, and genetic correlation estimates biased downwards.

### Winner’s curse is a function of architecture and power

Where the most statistically extreme associations from a GWAS are of interest to a follow up study, for example in MR, there is a potentially biasing influence of winner’s curse on the estimated effect sizes ^35^. Overcoming this issue can be achieved with the availability of independent replication of these effects. Unfortunately this is a challenge in the current data sharing ecosystem as the generation of replicated effect sizes is relatively ad hoc, and their dissemination is much harder to harvest than the discovery GWAS summary data. One advantage of OpenGWAS is that many traits have multiple datasets from independent samples, often owing to systematic analysis of traits in UK Biobank, so users can perform *post hoc* discovery and replication analyses within the database. A detailed exploration of one context in which the winner’s curse is a potentially serious issue - that of performing MR with weak instruments - is provided in Elsworth et al (2020), highlighting the importance of creating a systematic way of linking discovery and replication data in resources such as OpenGWAS.

### Summary

We have developed a system for housing GWAS summary data that aims to be scalable and programmatically accessible, while ensuring a level of data QC and harmonisation that simplifies downstream analyses.

## Web links

OpenGWAS homepage: https://gwas.mrcieu.ac.uk/

OpenGWAS API: https://gwas-api.mrcieu.ac.uk/

gwasglue R package: https://mrcieu.github.io/gwasglue/

gwasvcf R package: https://mrcieu.github.io/gwasvcf/

pygwasvcf python package: https://mrcieu.github.io/pygwasvcf/

ieugwasr R package: https://mrcieu.github.io/ieugwasr/

ieugwaspy python package: https://mrcieu.github.io/ieugwaspy/

TwoSampleMR R package: https://mrcieu.github.io/TwoSampleMR/

gwas2vcf command line tool: https://github.com/mrcieu/gwas2vcf

Data requests for OpenGWAS: https://github.com/mrcieu/opengwas-requests

## Acknowledgements

GH is funded by the Wellcome Trust and Royal Society [208806/Z/17/Z]. PCH was supported by CRUK Population Research Postdoctoral Fellowship C52724/A20138. Development of the platform was supported by a Wellcome Trust Institutional Translational Partnership Award (209739/Z/17/Z). The work was carried out in the UK Medical Research Council Integrative Epidemiology Unit (MC_UU_00011/1, MC_UU_00011/4). The project also received support from a Cancer Research UK programme grant (C18281/A19169), a British Heart Foundation Accelerator Award (AA/18/7/34219) and the NIHR Biomedical Research Centre at the University Hospitals Bristol NHS Foundation Trust and the University of Bristol.

## Competing interests

TRG, GH and GDS have received research funding from GlaxoSmithKline and Biogen for projects that use the MRC IEU OpenGWAS database. VH has previously been supported by funding from GlaxoSmithKline. Neither company had any input into or control over the contents of this manuscript.

Oracle have provided cloud resources to host the OpenGWAS database.

## Methods

### Obtaining GWAS summary data

GWAS summary data was obtained from several sources. Table 1 provides an overview of the data sources present in OpenGWAS at the time of writing. In general, we group datasets using the following rules:

**Table 1:** Data batches within the OpenGWAS

| **count** | **description** | **id** |
| --- | --- | --- |
| 293 | Datasets that satisfy minimum requirements imported from the EBI database of complete GWAS summary data<sup>15</sup> | ebi-a |
| 19942 | eQTLGen 2019 results, comprising all cis and some trans regions of gene expression in whole blood<sup>40</sup> | eqtl-a |
| 465 | GWAS summary datasets generated by many different consortia that have been manually collected and curated, initially developed for MR-Base<sup>6</sup> | ieu-a |
| 452 | Human blood metabolites analysed by Shin et al 2014<sup>41</sup> | met-a |
| 150 | Human immune system traits analysed by Roederer et al 2015<sup>42</sup> | met-b |
| 123 | Circulating metabolites analysed by Kettunen et al 2016<sup>43</sup> | met-c |
| 3282 | Complete GWAS summary data on protein levels as described by Sun et al 2018<sup>44</sup> | prot-a |
| 83 | Complete GWAS summary data on protein levels as described by Folkersen et al 2017<sup>45</sup> | prot-b |
| 3143 | Complete GWAS summary data on brain region volumes as described by Elliott et al 2018<sup>46</sup> | ubm-a |
| 2514 | IEU analysis of UK Biobank phenotypes<sup>47</sup> | ukb-b |
| 904 | Neale lab analysis of UK Biobank phenotypes, round 2<sup>48</sup> | ukb-d |

1. If one or a few summary datasets arise from a publication, we add these to the general data batch known as ‘ieu-a’. This contains a diverse collection of such published GWAS datasets collected over the last few years.
2. If there are a large number of summary datasets arising from a publication (e.g. a set of metabolites, protein levels, expression levels etc) or a systematic biobank-wide GWAS analysis that were generated using a single analytical pipeline, a new dedicated data-batch is created for that entire set of datasets (e.g. ukb-a, ukb-b, met-a, etc).

We also incorporate all harmonised datasets from the EBI GWAS summary database into OpenGWAS. The publicly available data can be searched and browsed online.

Our recommendation for users wishing to upload single published datasets is to deposit them in the EBI-NHGRI GWAS catalog, following which they will automatically appear in OpenGWAS after they have been through the EBI-NHGRI GWAS harmonisation pipeline.

For new data batches comprising large-scale systematic or unpublished datasets, users can submit a request for upload using a specific github issues page created for this process (https://github.com/mrcieu/opengwas-requests).

### Linkage disequilibrium reference panel

For some operations, such as linkage disequilibrium (LD) clumping or generating LD matrices for a set of variants, it is necessary to calculate the LD between a set of variants on the fly. For LD clumping operations we use five different super-populations from the 1000 genomes phase 3 sequenced reference panel, depending on the geographic origin of the individuals in the GWAS study. We retained only autosomal single nucleotide polymorphisms with minor allele frequency > 0.01 within the super-population. INDELs are removed, while multiallelic variants (with duplicate rsids representing the different allele pairs) are reduced to a single variant representing the most common allele pair.

The clumping endpoint of the API can be used to perform clumping using PLINK 1.90 on one of the five LD reference panels for a set of GWAS summary data. Very strict clumping thresholds are used by default - retaining variants with LD *R*^2^ < 0.001 within a 10Mb window and with p-value < 5*e*^−8^. While this is conservative, it ensures independence of top hits, but users can optionally implement alternative thresholds through the API.

The API is also able to provide a LD matrix utilising a similar approach in which the signed LD r value for the pairwise set of variants is returned as a matrix by issuing a PLINK 1.90 call to process the relevant variants using the specified LD reference panel.

### Data processing pipeline

Meta-data are stored as a flat file in JSON format, describing the GWAS file columns and the study metadata. Summary statistics are processed either on single datasets through the API, or for large numbers of datasets simultaneously in a HPC environment. The API orchestrates a containerised pipeline using the Cromwell scientific workflow engine (https://cromwell.readthedocs.io/en/stable/). The HPC pipeline orchestrates the same software natively but using a Snakemake (https://snakemake.readthedocs.io/en/stable/) pipeline.

GWAS summary statistics and study metadata are processed using gwas2vcf (https://github.com/mrcieu/gwas2vcf; Figure 2). First, the data are read and validated to ensure no missing values of essentials inputs and to ensure variables are of the correct type. Each variant is compared against a supplied reference sequence (GRCh37/hg19 by default), if the data is not already on the forward strand then the alleles are flipped to the forward strand where necessary. During this process the data is checked for errors, if each column of the data has the appropriate characteristics. For example, standard errors must be positive and numeric, effect sizes must be numeric, p-values must be between 0 and 1. All variants are oriented to the human genome reference panel (GRCh37/hg19) based on their chromosome and position, which means that the non-effect allele is forced to be the human genome reference allele. If this requires switching the effect allele, then the sign of the effect size is also switched and the reported effect allele frequency is transformed appropriately. Variants are subsequently left-aligned and trimmed to the minimal representation using vgraph (https://github.com/bioinformed/vgraph). This process ensures equal comparison of haplotypes between studies. The data are then annotated by position with the dbSNP database (build 153). Therefore, across all datasets every variant has a consistent rsID and effect alleles are consistently the same.

### Effect allele checks

An ever present concern with GWAS summary data in the current climate is that the effect allele has been incorrectly annotated, which could lead to analytical problems. For example, in a MR analysis the direction of a causal effect would be erroneously inferred to be the reverse. All studies in the OpenGWAS database have been verified for their effect allele frequency via documentation released with the original data or through personal communication. However we have also implemented a separate automatic check. Here, if the effect allele frequency is available then we compare the reported effect allele frequency for the study against a relevant population reference, using a set of common variants in the frequency range of 0.1-0.3. An inverse relationship between the effect allele frequency and the population allele frequency indicates the effect allele is incorrectly annotated, or the effect allele frequency is incorrectly annotated. Several studies have been flagged (across many of the batches) that have required follow up investigation to assert the correct allele frequency. This problem ambiguous GWAS summary data raises the importance of using standard GWAS summary data formats^23^.

### Per-study reports

Upon data conversion, a report is produced for visual inspection of the data. This involves performing LD score regression and LD-based clumping as described above. The Z-values obtained by dividing the effect size by the standard error are plotted against the Z-values obtained from the p-values to ensure that these data fields have been correctly specified. Effect allele frequencies are compared against the 1000 genome reference panel to help evaluate if chromosome positions and rsIDs have been specified correctly. Numbers of independent top hits are displayed, along with the estimates of genomic inflation factor and LD score regression outputs, to help evaluate if population structure is having a large influence on the traits and whether the number of independent top hits is reflective of the sample size of the data.

### LD score regression procedure

As part of the GWAS QC report, LD score regression was performed using the LDSC python package^36^ using default LD scores from European samples in the 1000 genomes reference panel. For each dataset variants were only retained if they were present in the hapmap3 reference panel^37^, excluding the MHC region.

### Database design

Once the QC report for a dataset has been inspected, an API call prepares the data for upload to be made public.

The data infrastructure has three main components: an Elasticsearch cluster (https://www.elastic.co/elasticsearch) to facilitate efficient search of the GWAS summary data, a Neo4j graph database (https://neo4j.com/) to store metadata for each study, and flat files that provide LD reference panels. The summary data and meta-data are all stored separately as flat files also, which can be downloaded directly in GWAS-VCF format and serve as the source of data fed into the Elasticsearch and Neo4j databases.

### Metadata

The metadata for each GWAS are stored in a unique node in the Neo4j graph, a description of which can be seen in table 2. In addition, information about GWAS access and permissions for embargoed datasets are also stored in the graph. Each database query is achieved via the API. For each query, the user ID is used to retrieve a list of available GWAS IDs from the Neo4j graph, which are then used to query the Elasticsearch cluster. Where possible, studies are mapped to existing ontologies, for example the trait name is mapped to an experimental factor ontology term (EFO) ^38^ and sample ancestry is mapped to the human ancestry ontology ^39^.

**Table 2:** GWAS metadata fields

| **Property** | **Description** |
| --- | --- |
| id | A unique ID |
| trait | Description of measured trait |
| note | Extra information |
| year | Year of publication |
| consortium | GWAS consortium name, if applicable |
| author | First author of GWAS |
| pmid | PubMed identifier, where available |
| sample_size | Sample size of GWAS |
| ncases | Number of cases |
| nsnp | Number of variants in results file |
| sex | Males/Females |
| population | Geographic origins of the samples included in the study |
| sd | Standard deviation of the phenotype analysed |
| unit | How to interpret a 1-unit change in the phenotype? eg log odds ratio, mmol/L, SD units |
| category | Binary disease phenotype or a non-disease phenotype |
| subcategory | Further categorisation |
| efo | List of experimental factor ontology terms |
| mr | Suitability of GWAS for Mendelian Randomization |

### GWAS summary data

The harmonised GWAS-VCF format summary data are served to the public from an Elasticsearch cluster, which is only accessible through the API. The Elasticsearch architecture was selected over other database platforms on the basis of query speed and scalability. The GWAS summary data are pre-indexed as part of the data processing pipeline, and these indexed data snapshots are deployed on an Elasticsearch cluster hosted on the Oracle Cloud Infrastructure (https://www.oracle.com/uk/cloud/).

Prior to indexing into Elasticsearch, all data were cleaned to remove non-numeric fields and missing values replaced with null and transformed into a set format (Table 3)

**Table 3:** GWAS summary data fields stored in the Elasticsearch database

| **Property** | **Description** |
| --- | --- |
| gwas_id | GWAS ID |
| chromosome | Chromosome hg19 |
| position | Position hg19 |
| effect_allele | Effect allele |
| other_allele | Alternative allele |
| effect_allele_freq | Effect allele frequency |
| p | P-value |
| n | Sample size for that variant |
| beta | Beta coefficient |
| se | Standard error |

**Table 4.**
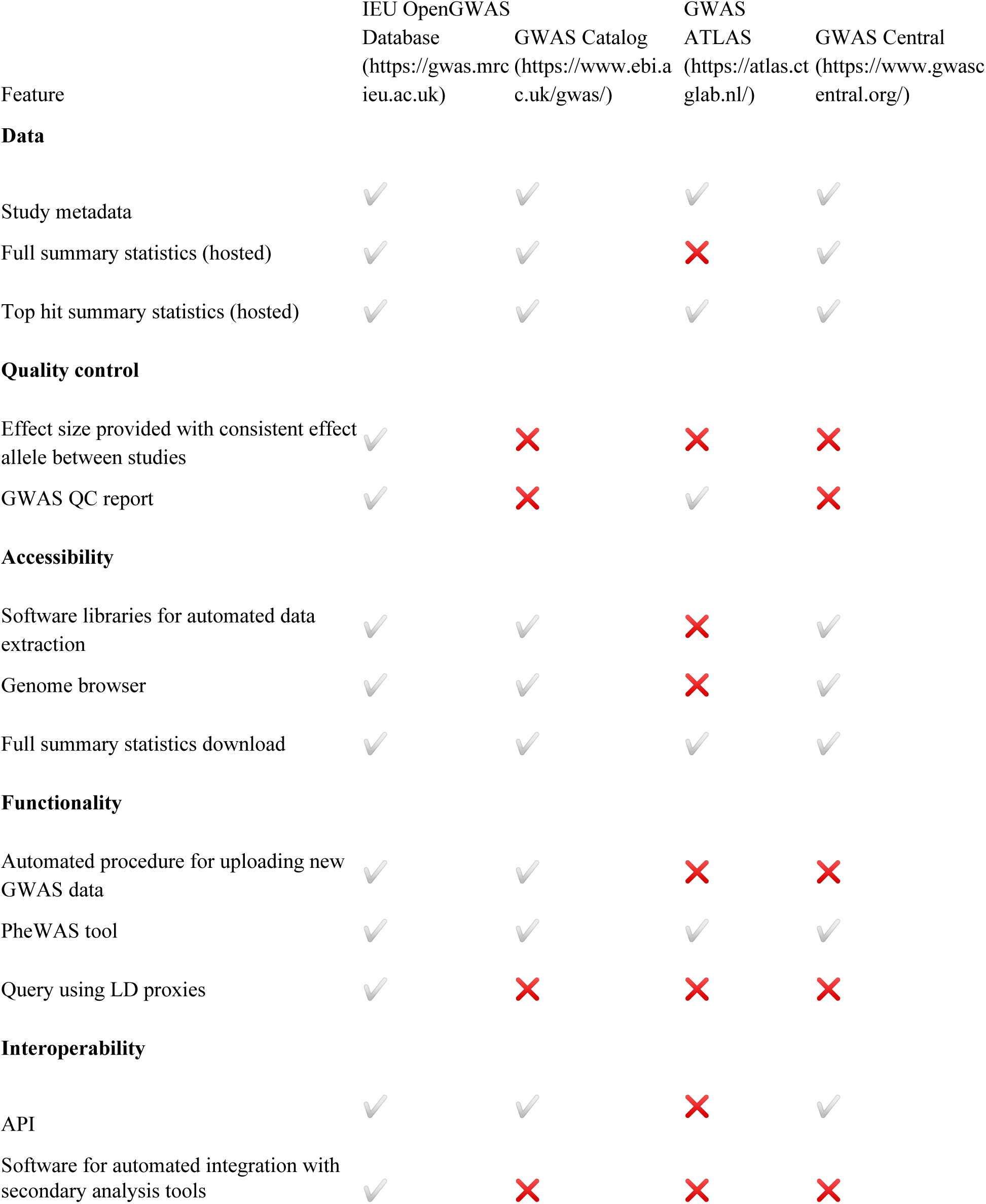
Comparison of IEU OpenGWAS infrastructure with existing tools

### API design

An application programming interface (API) is available for automated access to GWAS results and corresponding metadata. The Python3 API is built using Flask-RESTPlus, adhering to the representational state transfer (REST) architectural constraints. The swagger (https://swagger.io/) implementation utilizes the popular Flask framework with comprehensive unit and integration testing for accuracy and robustness. High availability and performance are achieved by running multiple containerised instances of the API behind a load balancing reverse proxy.

### LD proxy lookups

Pairwise LD *R*^2^ values are pre-calculated for variants within 10Mb windows, and pairs of variants are retained where LD *R*^2^ > 0.4. This list of pairs of variants, along with their LD *R*^2^ values and the allelic phase of the two variants, are stored within the Elasticsearch database. If a variant is requested for a dataset and is found not to be present then the following procedure occurs:

1. The LD pair database is searched, identifying the variants with highest LD with the target variant.
2. The target dataset is queried for each of the proxy variants from (1).
3. The proxy variant present in the target dataset with the highest *R*^2^ with the target variant is retained.
4. The allele of the proxy variant that is in phase with the effect allele of the target variant is identified.
5. The effect, with respect to the proxy effect allele from (4), is returned as the proxy effect for the target variant.

